# Lysyl oxidase promotes neuronal ferroptosis exacerbating seizure-induced hippocampal damage

**DOI:** 10.1101/839852

**Authors:** Xiao-Yuan Mao, Ming-Zhu Jin, Qin Li, Ji-Ning Jia, Qian-Yi Sun, Hong-Hao Zhou, Zhao-Qian Liu, Wei-Lin Jin

## Abstract

Epilepsy is a serious neurological disorder and characterized by recurrent and unprovoked seizures. A critical pathological factor in the seizure genesis is neuronal loss. However, mechanisms which lead to neuronal death remain elusive. Our present investigation depicted that ferroptosis, a recently discovered iron- and lipid peroxidation-dependent cell death, probably served as a mechanism in murine models of kainic acid (KA)-induced seizures. And treatment with ferroptosis inhibitors ferrostatin-1 (Fer-1), liproxstatin-1 (Lipo-1) or deferoxamine (DFO) significantly suppressed seizure severity and frequency. Using gene expression profiling in HT22 cells after glutamate exposure (a validated ferroptotic cell death model), we identified lysyl oxidase (Lox) as a novel inducer of ferroptosis. Mechanistically, Lox promoted ferroptosis via activation of extracellular regulated protein kinase (ERK)-dependent 5-lipoxygenase (Alox5) phosphorylation at serine 663 residue signaling, subsequent leading to lipid reactive oxygen species (ROS) accumulation. In a murine model of KA-induced seizure, we illustrated that administration of β-aminopropionitrile (BAPN), a specific Lox inhibitor, remarkably prevented seizure generation. Overall, these findings highlight Lox, a novel identified ferroptotic regulator in neurons, serves as a potential target for seizure-related disease including epilepsy.

## Introduction

Epilepsy is by far the most prevalent serious neurological disorder worldwide, affecting nearly 50 million people. It is often featured by an enduring propensity for generation of unprovoked epileptic seizures in the vulnerable brain area including hippocampus (1, 2) and the associated neurological comorbidities such as cognitive deterioration, psychological and social consequences, finally seriously affecting the life quality of epileptic patients. Seizure is a common phenomenon in the etiology of epilepsy and it is well known that severe and repetitive seizures can result in activation of neuronal death pathways (2). In fact, hippocampal neuronal loss always occurs in epileptogenic foci (3). However, the potential molecular mechanism of neuronal loss remains enigmatic. It was previously unequivocally reported that seizure-activated neuronal cell death pathway could lead to progressive hippocampal neuronal loss and cognition deficits (4). Deciphering the molecular pathways underlying seizure-induced neuronal death is invaluable to the development of novel therapeutic approaches for protecting the brain against prolonged or repetitive seizures during epileptogenesis.

Classically, neuronal death modality has been divided into two major forms: apoptosis and necrosis, on the basis of their distinct morphological, biochemical and molecular attributes. Apoptosis is a sort of programmed cell death while necrosis is considered to be uncontrolled (5). More recently, there is now evidence supporting that necrosis can be regulated by genetic and pharmacological ways, which is called “regulated necrosis” (6-8). Nowadays, multiple paradigms of regulated necrosis process have been discovered, including necroptosis, pyroptosis and ferroptosis (9-11).

Ferroptosis is a recently discovered form of regulated cell death (RCD) which differs from other cell death modes by its characteristics of mitochondrial shrinkage and condensed membrane density (morphological), iron-dependent lethal lipid-based reactive oxygen species (ROS) overproduction, particularly lipid hydroperoxide (biochemical) and involvement of unique execution mechanisms (11). Lipid peroxidation is a critical feature in the ferroptosis execution. There is ample evidence supporting that various modulators of lipid metabolism including Acyl-CoA synthetase long-chain family 4 (ACSL4), glutathione peroxidase 4 (GPX4), 5-lipoxygenase (Alox5), Alox12 and phosphatidyletha-nolamine-binding protein 1 (PEBP1) are involved in ferroptotic cell death (12-16). For instance, it has demonstrated that an enhanced Alox5 activity has been found in ferroptosis process under glutamate-triggered neurotoxicity in hippocampal neurons and pharmacological inhibition of Alox5 by Zileuton counteracts ferroptotic death event in this model (14). Nowadays, ferroptosis has been extensively reported in multiple pathogenic conditions including cancers (17-19), neurological disorders (20, 21) and renal failure (13). Particularly, our recent publications (22, 23) have unequivocally proved the occurrence of ferroptosis in various chemical reagents-induced epileptic rodent models. And treatment with ferroptosis inhibitors (ferrostatin-1 (Fer-1) or liproxstatin-1 (Lipo-1)) efficiently alleviates seizures, highlighting a crucial role of ferroptosis process in seizure generation. In this respect, a deep understanding of the potential mechanisms that regulate ferroptosis might provide alternative therapeutic targets for treating seizure-associated disorders including epilepsy.

In our present work, it is revealed that ferroptosis is present in a mouse model of kainic acid (KA)-induced seizures and human epilepsy. And treatment with ferroptosis inhibitors Fer-1, Lipo-1 or Deferoxamine (DFO) evidently alleviates epileptic seizures in mice. Furthermore, using gene expression profile, we identify and validate lysyl oxidase (Lox), a copper-dependent monoamine enzyme in the extracellular matrix (ECM) remodeling, as a novel ferroptotic inducer in vitro hippocampal HT22 neuron ferroptosis models caused by glutamate or erastin. Pharmacological inhibition of Lox by a specific inhibitor β-aminopropionitrile (BAPN) prominently ameliorates epileptic seizures.

Lox is initially responsible for the catalysis of cross-linking of elastin and collagens in the ECM (24). Previous investigations have indicated its contribution to tumor metastasis and pathogenesis of hypertension (25, 26). Nevertheless, little is known of the roles of Lox in seizure genesis and/or the progression of epilepsy. Our present investigation deciphers Lox promotes neuronal ferroptosis via activating ERK-Alox5 signaling in epileptic seizures and this work may provide us new insights into clinical translation of the axis as an early target for therapeutic intervention of epileptic seizures.

## Results

### Ferroptosis is activated in KA-induced seizure mice and human epileptic patients

Our prior work showed the presence of ferroptosis in various epileptic seizure models induced by chemical reagents including pentylenetetrazol, pilocarpine and FeCl_3_ (22, 23). To determine whether ferroptosis process was activated in KA-induced seizures, a common model resembling human intractable epilepsy, we detected a set of ferroptotic indices in the hippocampus of this model including morphological characteristics of mitochondrion, lipid peroxidation by-products including 4-HNE and MDA levels and *Ptgs2* mRNA. It was noteworthy that smaller and more condensed mitochondria (Figure 1A) and elevated levels of 4-HNE (Figure 1B and Figure 1C), MDA (Figure 1D) and *Ptgs2* mRNA (Figure 1E) were found in mice subjected to KA-induced seizure, which fit the major criteria for ferroptosis. Besides, *Ptgs2* mRNA level was also observed to be significantly augmented in brain tissue samples of human epilepsy compared with non-epileptic patients (Figure 1F). Collectively, these findings implicate that ferroptosis occurs in a mouse model of KA-induced seizures and epileptic patients.

**Figure 1.**
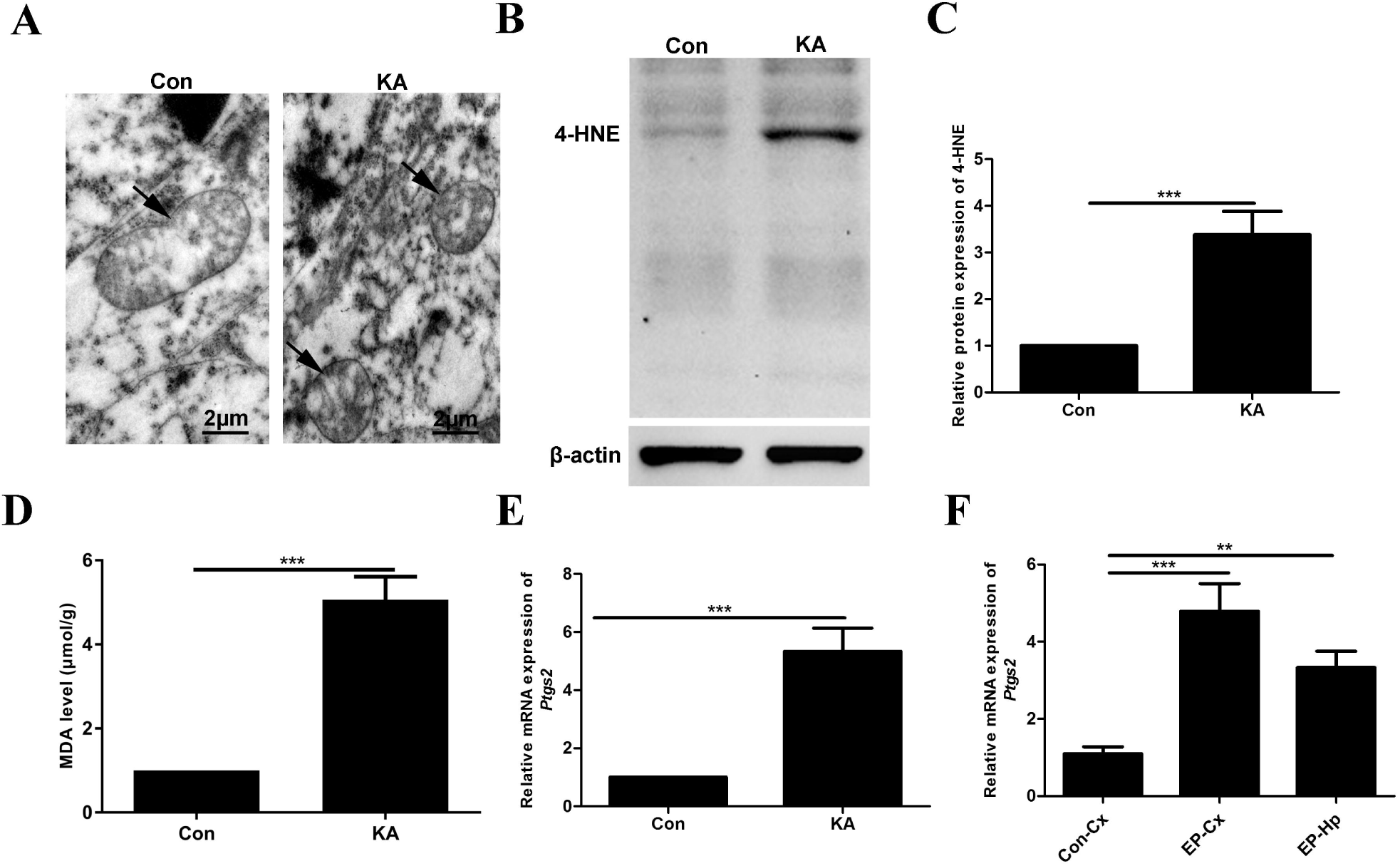
Ferroptosis activation in KA-induced seizures in mice and human epileptic patients. **(A)** Transmission electron microscopy of brain tissue samples for analyzing morphological alteration of mitochondria. Arrows indicate mitochondria. Scale bar: 2 μm. **(B-C)** Protein expression of 4-HNE by Western blot. **(D)** Measurement of MDA level by a commercial assay kit. **(E-F)** Determination of *Ptgs2* mRNA by quantitative real-time PCR in KA-induced seizures in mice and human epileptic patients. n=6 for each group, \*\**p*<0.01 and \*\*\**p*<0.001.

### Suppression of ferroptosis process improves phenotype in a mouse model of KA-induced seizures

Next, to investigate the contributions of in seizures, we evaluated seizure phenotype in mice subjected to KA via seizure latency, seizure score and number of seizures within 90 min after treatment with ferroptosis inhibitors including Fer-1, Lipo-1 or DFO (Figure 2A). Intriguingly, it was observed that Fer-1, Lipo or DFO all remarkably ameliorated seizure behavior (Figure 2B and *Supplementary Video*). In details, seizure score (Figure 2D) and number of seizures within 90 min (Figure 2E) were both prominently decreased in KA-triggered seizures in mice when treatment with Fer-1, Lipo-1 or DFO, despite no significant difference for seizure latency (Figure 2C). These data suggest that blockade of ferroptosis has potential therapeutic value for seizure control.

**Figure 2.**
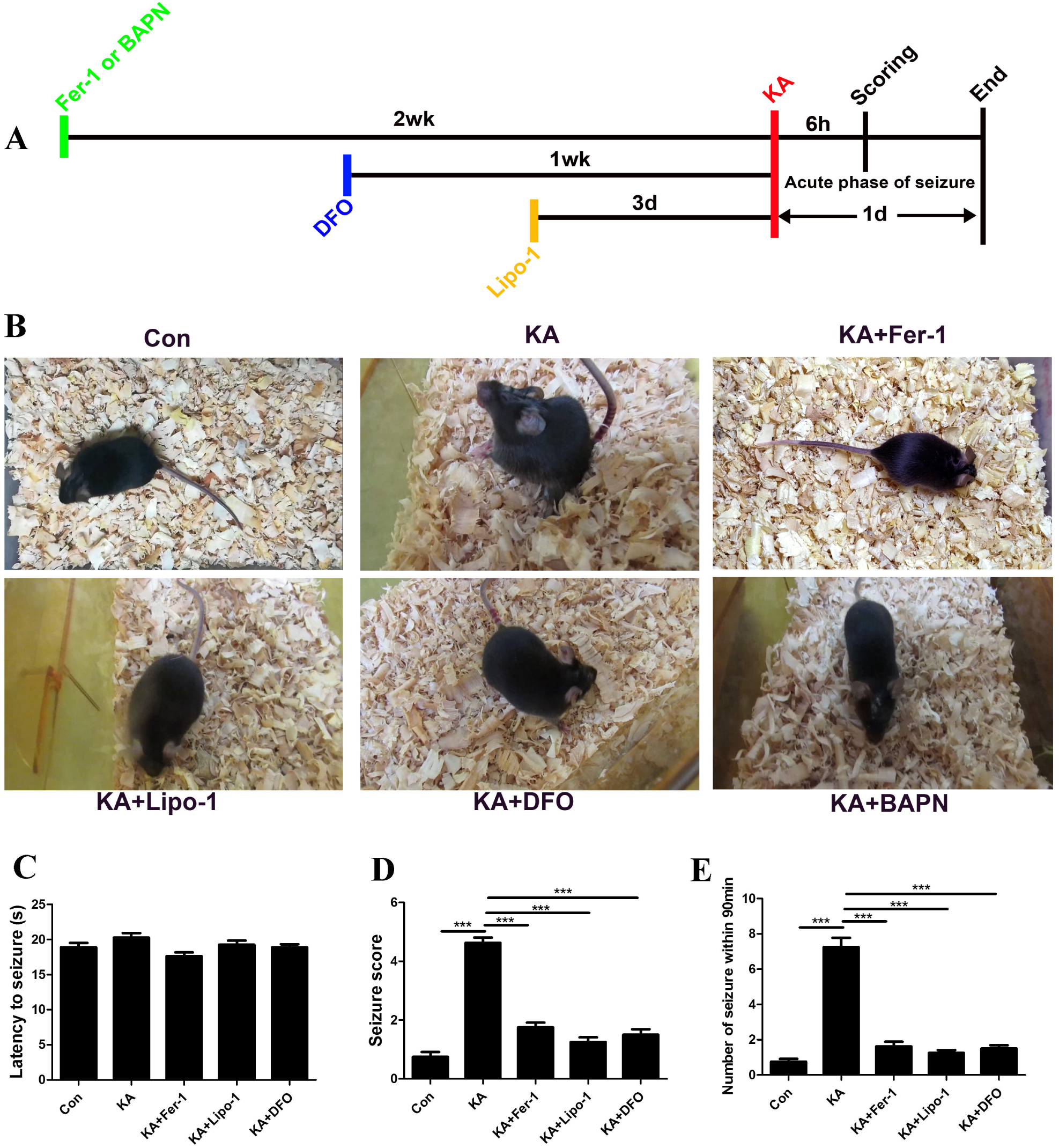
Inhibition of ferroptosis process ameliorates seizures in KA-treated mice. **(A)** Experimental procedure. **(B)** Representative images showing improvement of ferroptosis inhibitors including Fer-1, Lipo-1 and DFO and Lox inhibitor BAPN on seizure behavior. **(C-E)** Effects of Fer-1, Lipo-1 and DFO on latency to seizure, seizure score and number of seizures. n=8 for each group, \*\*\**p*<0.001.

### Lox functions as a novel ferroptosis contributor in glutamate-induced oxidative toxicity in hippocampal HT22 neurons

To further probe the potential molecular mechanism underling the involvement of ferroptosis in seizures, we firstly established a model with glutamate-induced oxidative toxicity in the immortalized HT22 hippocampal neurons. HT22 has the feature of deficient ionotropic glutamate receptors (27) and has been considered to be an excellent model for the study of glutamate-mediated oxidative impairment. As shown in Figure 3A, multiple cell death modalities including ferroptosis, autophagy and necroptosis existed in glutamate-stimulated oxidative stress in HT22 cells as pretreatment with the corresponding cell death inhibitors (Fer-1, Lipo-1 and DFO for blocking ferroptosis, 3-MA for inhibiting autophagy and Nec-1 for suppressing necroptosis) all protected neuronal HT22 cells from oxidative damage. What was more important, pre-incubation with specific ferroptosis inhibitors Fer-1 (11), Lipo-1 (13) or DFO (11) constituted the most effective intervention in comparison to treatment with autophagic cell death suppressor 3-MA or necroptosis inhibitor Nec-1, in glutamate-induced HT22 oxidative toxicity. To further validate the existence of ferroptosis in this model, ferroptotic indices including lipid ROS (Figure 3B), 4-HNE protein level (Figure 3C), MDA content (Figure 3D) and *Ptgs2* mRNA (Figure 3E) were also detected in HT22 cells following treatment with glutamate at different time points. In terms of lipid peroxidation, treatment with glutamate (5 mM) resulted in a time-dependent increase of lipid ROS and lipid degradation products (4-HNE and MDA), beginning at 4 h. This accumulation of lipid peroxidation preceded cell death, which began at 8 h (*Supplementary Figure S1*). In glutamate-induced oxidative toxicity in HT22 cells, ferroptotic cell death was also confirmed by the upregulation of *Ptgs2* at mRNA level, which was recently identified as a molecular biomarker of ferroptosis (28). Taken together, these findings indicated that ferroptosis served as a dominant cell death modality in glutamate toxicity in HT22 cells. In our subsequent work, this model was therefore employed to elucidate the molecular mechanisms leading to neuronal ferroptosis after glutamate-triggered oxidative stress in HT22 cells. Using gene expression profiles, we compared differentiated genes between glutamate (5 mM)- and vehicle-treated groups. Totally, 32 differentially expressed genes were found between these two groups. For details, 14 up-regulated genes and 18 down-regulated genes were observed in glutamate-treated group, compared with vehicle group (Figure 3F). In the validation stage, lysyl oxidase (Lox), a key enzyme for extracellular matrix (ECM) remodeling, was selected as a novel ferroptotic inducer since it was confirmed to be remarkably increased at different time points after glutamate exposure (Figure 3F) and prior study depicted that Lox induced oxidative stress in pathological scenarios (25). Additionally, evidence for the critical role of ECM in ferroptosis arise from the fact showing that α6β4 integrin-deficient cells are more vulnerable to ferroptosis (29). Lox protein level and Lox activity were further found to be enormously elevated in glutamate-treated group in a time-dependent manner (Figure 3G and Figure 3H). Consistently, enhanced mRNA and protein levels and activity of Lox were also verified in HT22 cells treated with erastin, a specific ferroptosis inducer (*Supplementary Figure S2*), while other Lox family members including Loxl1, Loxl3 and Loxl4 were all unchanged at transcriptional level either in glutamate- or erastin-induced neuronal ferroptosis (Loxl2 mRNA was undetectable) (*Supplementary Figure S3*). After knock down of Lox (Figure 3I) by genetic silencing, glutamate-induced cell death and lipid ROS were obviously impeded (Figure 3J and Figure 3K). Additionally, blockade of Lox by pharmacological inhibitor BAPN or binding to the active site with Lox antibody both obtained similar results as using genetic silencing (Figure 3L and Figure 3M). We also examined whether Lox was altered in other classical cell death modes including staurosporine-induced apoptosis and rapamycin-induced autophagy in HT22 neurons. It was noteworthy that no significant difference of Lox in apoptotic or autophagic cell model compared with control group (*Supplementary Figure S4*). These data suggest that Lox serves as a novel ferroptosis-inducing factor in glutamate-induced oxidative toxicity in hippocampal HT22 neurons.

**Figure 3.**
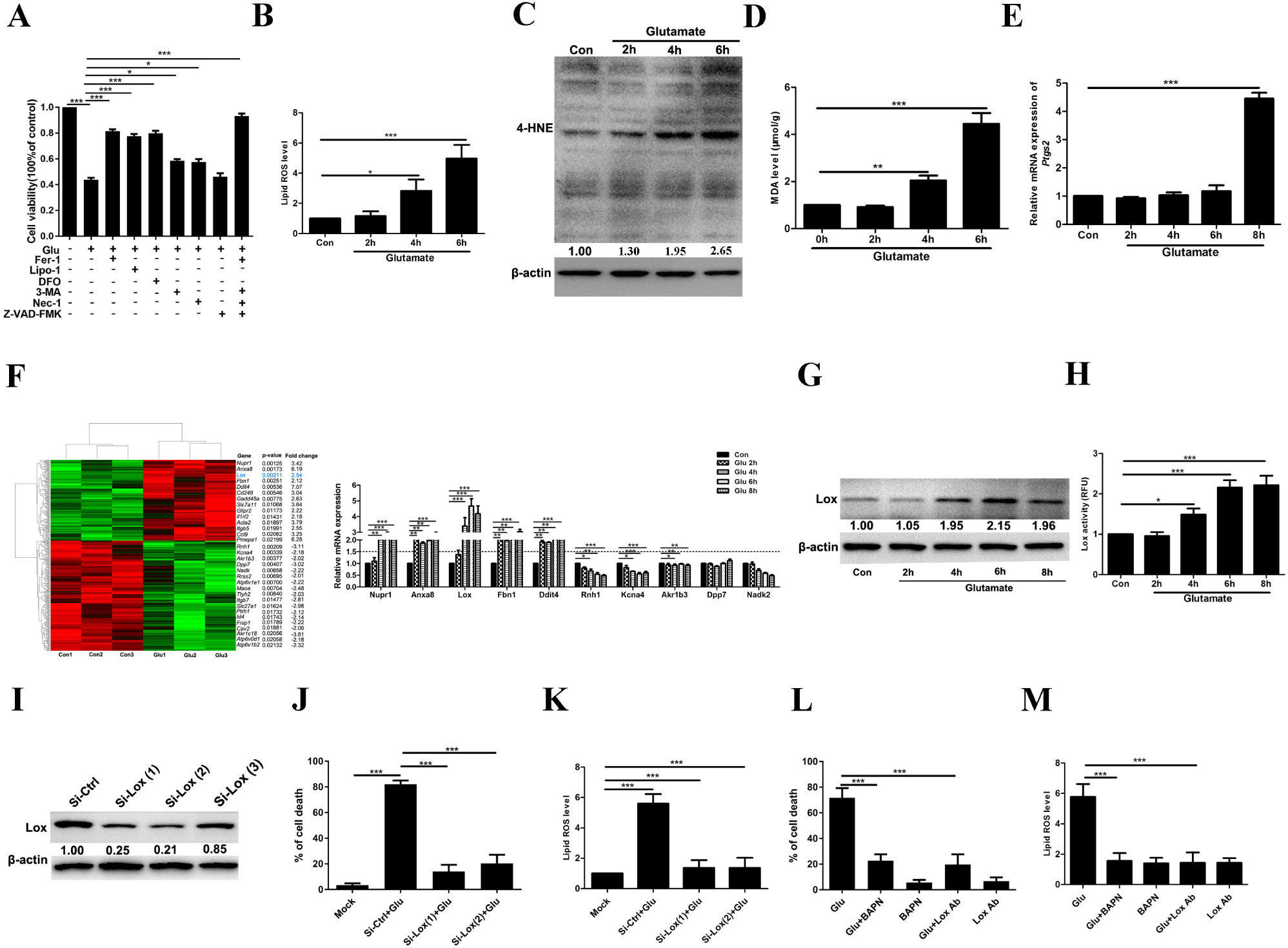
Lysyl oxidase serves as a novel ferroptotic regulator in HT22 neurons under glutamate exposure. **(A)** Cell viability of HT22 neurons pretreated with 12.5 μM Fer-1, 1 μM Lipo-1 and 50 μM DFO (ferroptosis inhibitors), 5 mM 3-MA (an autophagy inhibitor), 100 μM Nec-1 (a necroptosis inhibitor) or 100 μM Z-VAD-FMK (an apoptosis inhibitor) for 2 h, followed by 5 mM glutamate for 8 h. **(B)** Lipid ROS levels were measured in HT22 cells under glutamate stimulation at 2 h, 4 h and 6 h. **(C-E)** showed detections of 4-HNE, MDA and *Ptgs2* mRNA at different time points (2 h, 4 h and 6 h) under glutamate exposure, respectively. **(F)** Heat map showing differentially expressed genes in HT22 neurons between glutamate- and vehicle-treated groups. Down-regulated genes were indicated in green, and up-regulated genes were depicted in red. The top five up-regulated and the top five down-regulated gene expressions were further validated by quantitative real-time PCR. **(G)** Protein levels of Lox at different time points (2 h, 4 h, 6 h and 8 h) under glutamate exposure. **(H)** Determination of Lox activity at different time points (2 h, 4 h, 6 h and 8 h) under glutamate exposure. **(I)** Effects of Lox knockdown by genetic silence on Lox protein level. **(J-K)** displayed effects of Lox knockdown on cell survival and lipid ROS level, respectively; **(L-M)** showed effects of blockade of Lox by Lox antibody (Lox Ab) or a specific inhibitor, BAPN, on cell survival and lipid ROS level, respectively. \**p*<0.05, \*\**p*<0.01 and \*\*\**p*<0.001.

### Erk-Alox5 axis is a major downstream signaling for the contribution of Lox to ferroptosis in glutamate toxicity in hippocampal neurons

Next, we sought to decipher the mechanism of Lox-induced ferroptosis in oxidative glutamate toxicity in neurons. Figure 4A displayed the experimental procedures. Prior work showed Lox had the ability to enhance vascular oxidative stress via promoting p38 MAPK activation and subsequently facilitate elastin remodeling in hypertension (25). Our present investigation aimed to explore whether activated MAPK pathways are involved in ferroptosis induction by Lox. Under glutamate toxicity in HT22 neurons, Lox knock down via genetic silencing or pharmacological inhibition of Lox by BAPN both dramatically suppressed ERK phosphorylation while phosphorylations of JNK and p38 were unaltered (Figure 4B and Figure 4C). In contrast, suppression of ERK activation using a specific inhibitor U0126 was not able to alter Lox expression, but decreased Alox5 phosphorylation at serine 663 residue (Figure 4D). And in line with previous investigations (14, 30), ERK blockade by U0126 or inhibition of Alox5 by Zileuton obviously inhibited ferroptosis process and abrogated neuronal impairment induced by glutamate (Figure 4E and Figure 4F). Similarly, suppression of phosphorylated ERK by U0126 could also inhibited erastin-induced neuronal ferroptosis in HT22 cells while treatment with either a p38 protein kinase inhibitor (SB202190) or a JNK protein kinase inhibitor (SP600125) did not protect neurons from cell death (*Supplementary Figure S5*). Collectively, these results implicate that ERK-activated Alox5 ser663 phosphorylation functions as a downstream signaling pathway of Lox-mediated ferroptotic cell death.

**Figure 4.**
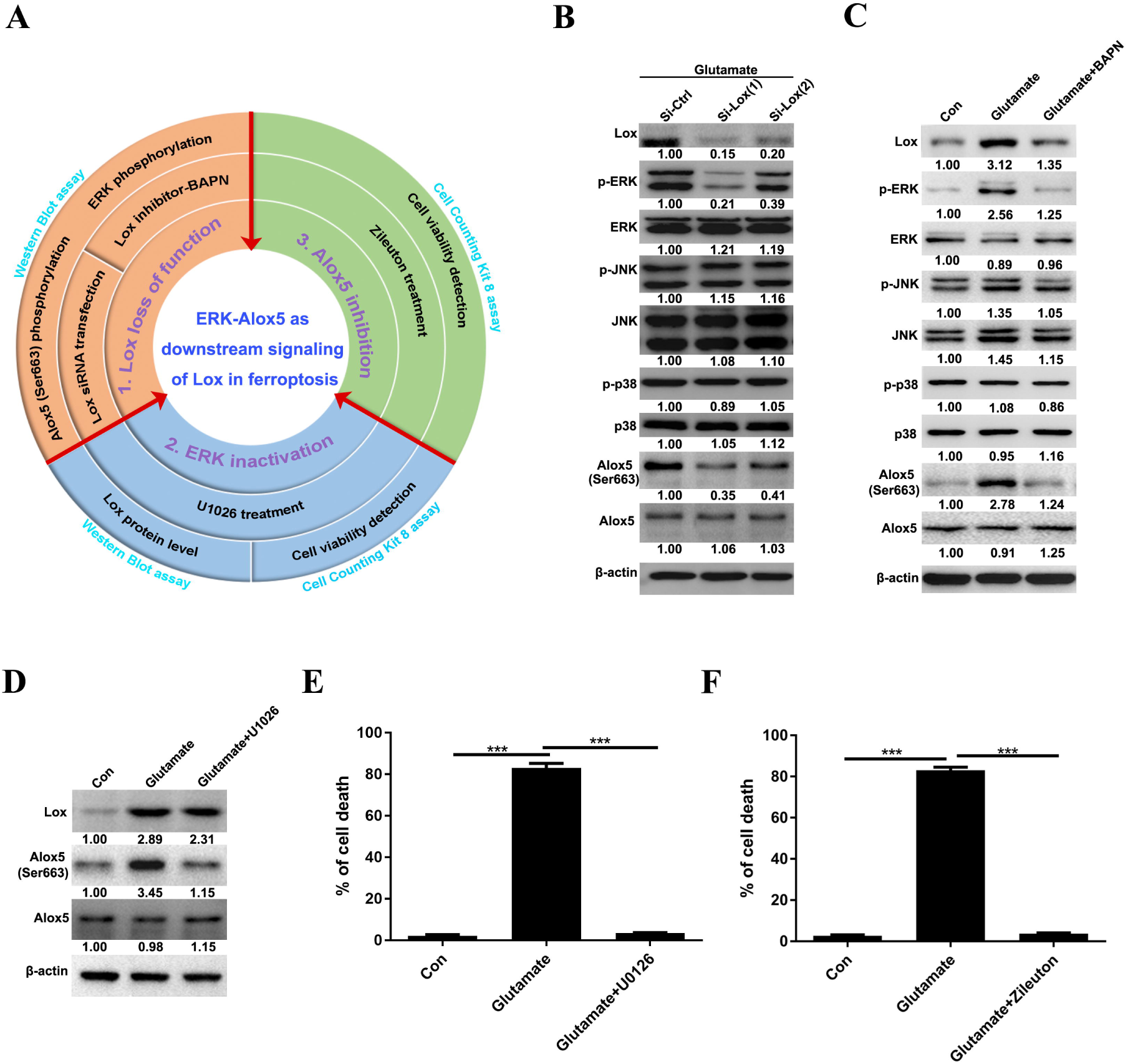
ERK-Alox5 signaling axis is involved in Lox-mediated ferroptotic process in HT22 neurons under glutamate exposure. **(A)** Experimental design. **(B)** Under glutamate toxicity, effects of Lox knockdown on MAPK pathway proteins (p-ERK, ERK, p-JNK, JNK, p-p38 and p38), Alox5 protein phosphorylation and total Alox5 protein levels. **(C)** Under glutamate toxicity, effects of BAPN on MAPK pathway proteins (p-ERK, ERK, p-JNK, JNK, p-p38 and p38), Alox5 protein phosphorylation and total Alox5 protein levels. **(D)** Under glutamate toxicity, effects of blockade of ERK by a specific inhibitor, U0126, on the protein levels of Lox, phosphorylated Alox5 and total Alox5. **(E-F)** showed effects of U0126 or Alox5 inhibitor (zileuton) on cell survival in glutamate-induced HT22 cell death, respectively. \*\*\**p*<0.001.

### Pharmacological inhibition of Lox by BAPN reduces seizure activity in mice

Subsequently, we explored whether inhibition of Lox could attenuate seizure phenotype in KA-treated mice. Expression levels of Lox mRNA and protein were confirmed to be evidently increased in the hippocampus of KA-induced epileptic seizure model (Figure 5A, Figure 5B and Figure 5C). We further validated that Lox was localized in neurons as evident overlapping was observed for Lox and NeuN, a specific neuronal marker (Figure 5D). In order to explore the role of Lox in the epileptic seizures, we investigated whether suppression of Lox by BAPN, a previously reported irreversible inhibitor (31), could impede these pathological conditions. The experimental procedure was displayed in Figure 2A. It was found that treatment with BAPN significantly suppressed seizure behavior in KA-treated mice (and *Supplementary Figure S1*). Further data analysis revealed that seizure score and the number of seizures were remarkably decreased in KA-induced epileptic seizures in mice after receiving different doses of BAPN (100 mg/kg or 300 mg/kg) evidently diminished (Figure 5F and Figure 5G), despite no significant difference in the aspect of latency to seizures (Figure 5E). Additionally, ferroptotic events were suppressed in this seizure model after BAPN treatment as reduced levels of *Ptgs2* mRNA and MDA level were observed (Figure 5H and Figure I).

**Figure 5.**
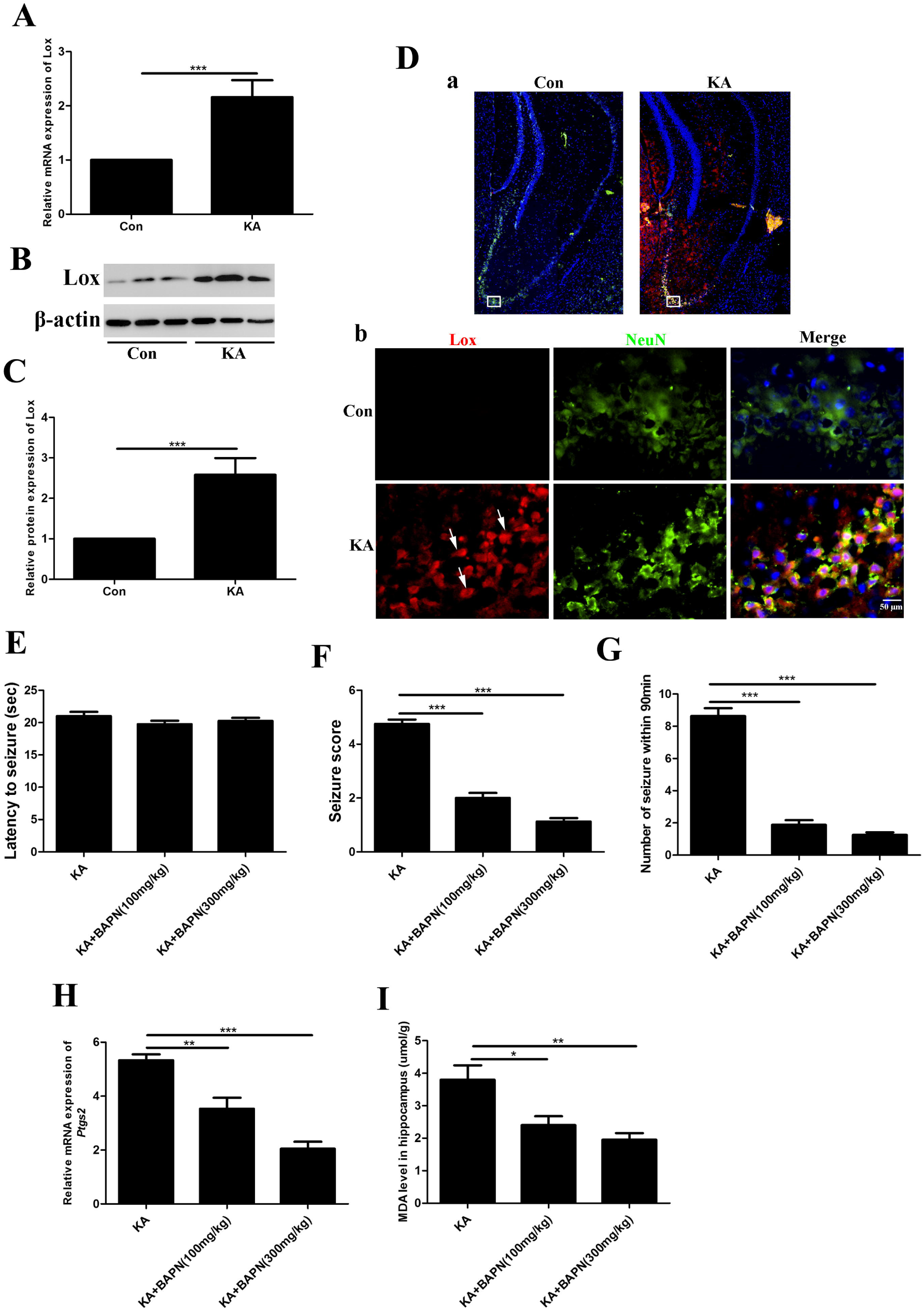
Lox inhibition by β-aminopropionitrile (BAPN) exerts anti-convulsant potential in vivo. **(A-C)** indicated Lox expression at mRNA and protein levels in a mouse model of KA-induced epileptic seizures and control group. **(D)** Immunofluorescence of Lox and NeuN in hippocampus of mice after KA-induced seizures and control group, Scale bar: 10 μm. **(E-G)** Effects of BAPN on seizure latency, seizure score and number of seizures within 90 min in a mouse model of KA-induced seizures. **(H-I)** showed effects of BAPN on ferroptotic induces including *Ptgs2* mRNA and MDA content in a mouse model of KA-induced seizures. \**p*<0.05, \*\**p*<0.01 and \*\*\**p*<0.001.

### AAV-mediated Lox transfer to hippocampal neurons exacerbates seizures and can be rescued by BAPN

Given the anti-seizure potential of Lox inhibitor, we further explored whether overexpression of Lox in neurons could aggravate seizures in KA-treated mice. Delivery of Lox in hippocampal neurons was performed via stereotactic injection of recombinant AAV-PHP.eB expressing Lox into hippocampus. The experimental protocol was listed in Figure 6A. The virus was confirmed to be successfully transferred into neurons as a strong overlap with GFP flag and NeuN via confocal analysis (Figure 6B). Lox overexpression was validated by Western blot analysis (Figure 6C). And compared with AAV-NC group, mice with infusion of recombinant AAV-Lox exhibited significantly increased seizure score and number of seizures in sub-convulsant dose of KA (Figure 6E and Figure 6F), indicating that Lox up-regulation in hippocampal neurons augments seizure sensitivity in mice. In order to further explore whether overexpression of Lox could also affect ferroptosis process in seizures, we detected ferroptotic indices including 4-HNE, MDA and *Ptgs2* (Figure 6G, Figure 6H, Figure 6I and Figure 6J). It was obviously found that recombinant Lox overexpression resulted in elevations of 4-HNE, MDA and *ptgs2* in KA-treated mice, compared with sub-convulsant dose (100 ng in 1μl saline) of KA-treated group. These findings again confirm that up-regulation of Lox promotes neuronal ferroptosis in seizures. In the meantime, the enhanced seizure sensitivity via AAV-mediated Lox transfer in the brain was rescued by pharmacological inhibition of Lox inhibitor BAPN, as shown in Figure 6L and Figure 6M, despite no difference in seizure latency (Figure 6K). Mechanistically, Lox overexpression in vivo contributed to increased phosphorylated protein levels of ERK and Alox5 (Ser663) (Figure 6N and Figure 6O). These data raised the possibility that up-regulation of Lox activated ERK-dependent Alox5 Ser663 phosphorylation signaling facilitated neuronal ferroptosis in seizure-induced hippocampal damage.

**Figure 6.**
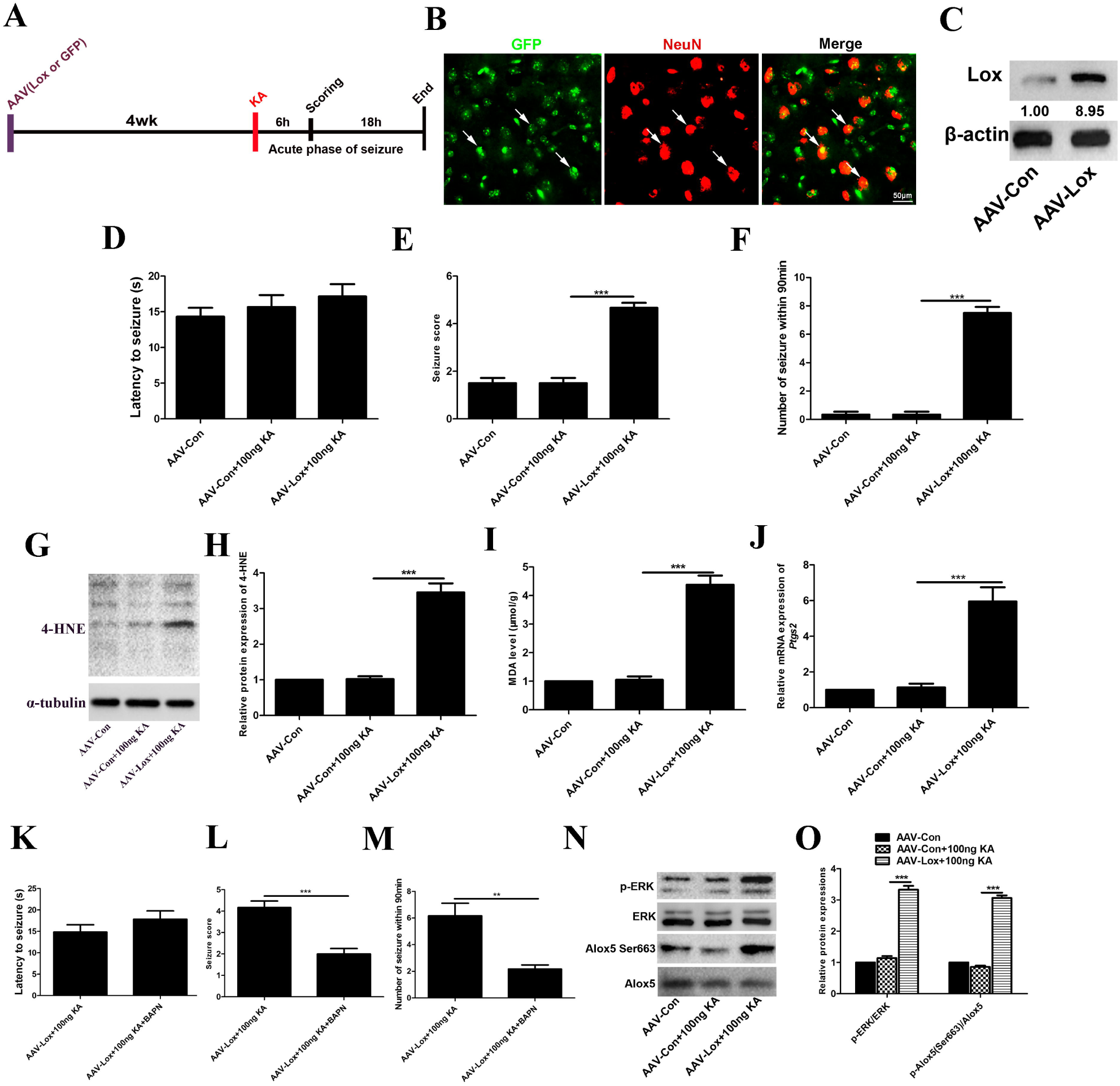
AAV-mediated Lox transfer to hippocampal neurons exacerbates seizures and can be rescued by BAPN. **(A)** Experimental workflow of AAV-mediated Lox transduction in vivo. **(B)** AAV-PHP.eB-GFP was injected into hippocampus to indicate neuron-specific expression. Representative images showed co-localization of GFP and NeuN in the AAV-injected hippocampus. Scale bar: 10 μm. **(C)** Validation of Lox overexpression in hippocampus by AAV-PHP.eB vector. **(D-F)** Evaluation of seizure severity and seizure frequency by seizure latency, seizure score and number of seizures within 90 min in mice subjected to a sub-convulsant dose of KA after AAV-mediated Lox transduction. **(G-J)** Effects of AAV-mediated Lox transduction on ferroptotic events (4-HNE, MDA and RNA expression of *Ptgs2*) in mice treated with a sub-convulsant dose of KA. **(K-M)** Effects of BAPN on seizure latency, seizure score and number of seizures within 90 min in mice treated with a sub-convulsant dose of KA after AAV-mediated Lox overexpression. **(N-O)** Effects of AAV-mediated Lox transduction on ERK phosphorylation and Alox5 Ser663 phosphorylation in mice treated with a sub-convulsant dose of KA. In the whole figure, 100 ng KA indicated injection of 100 ng KA in 1 μl saline. \*\**p*<0.01 and \*\*\**p*<0.001.

## Discussion

In the current work, we identify Lox as a novel neuronal ferroptosis inducer and mechanistically, ERK-Alox5 signaling serves as the downstream pathway of Lox to promote ferroptosis process in neurons. The Lox inhibitor BAPN can blunt ERK-Alox5-dependent ferroptosis process and exert potent anti-seizure potential (Figure 7B).

**Figure 7.**
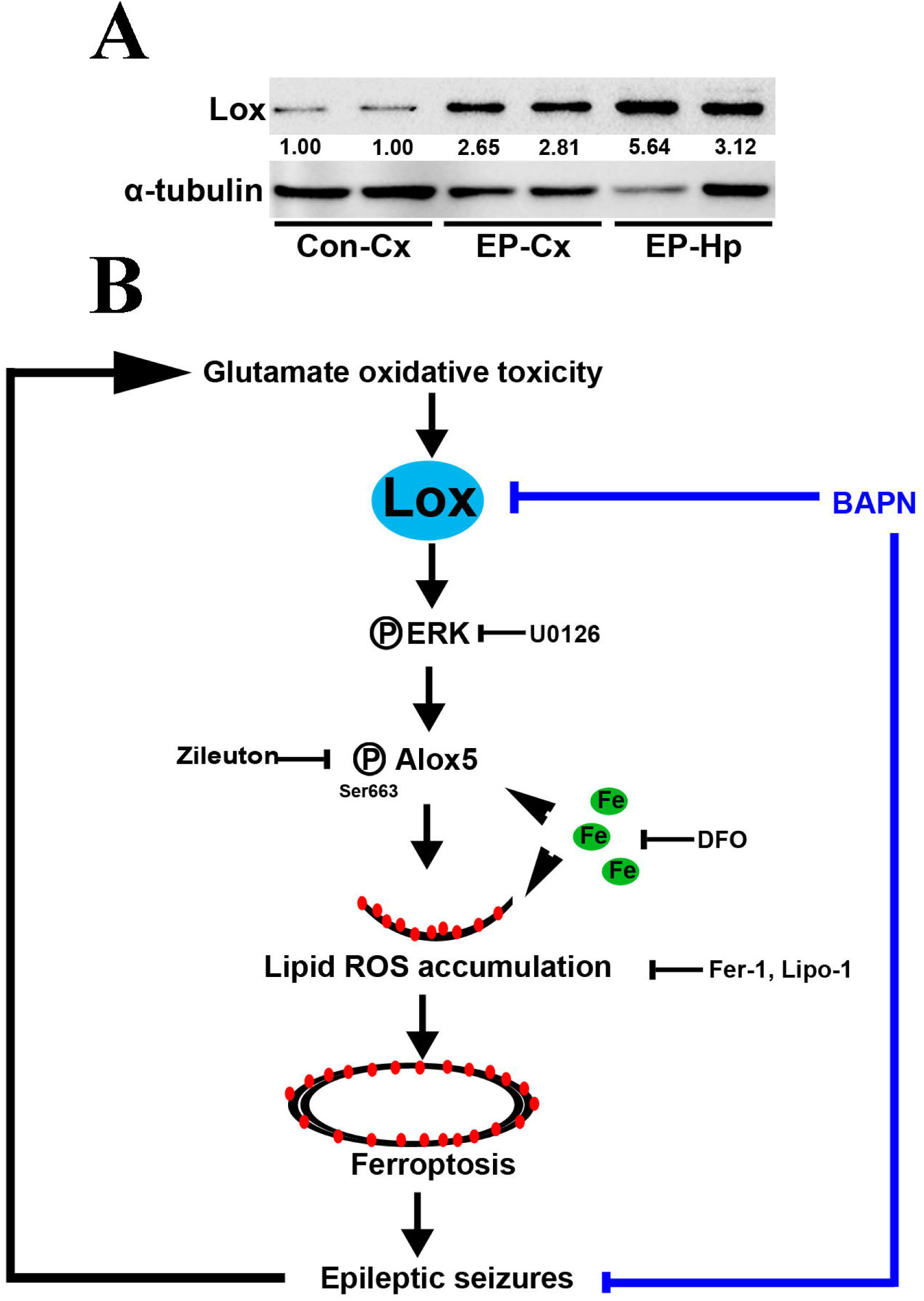
Working model. **(A)** Protein expression of Lox in brain tissues in human epileptic patients and non-epileptic control group. n=2 for each group. **(B)** Schematic representation of the role of the ERK-Alox5 signaling axis in glutamate oxidative toxicity-induced Lox expression and the role of Lox, a newly identified ferroptotic regulator in neurons, in seizure-induced hippocampal damage. Inhibition of Lox is a promising therapeutic approach for seizure-associated disorders such as epilepsy.

Seizures are very common in a plethora of human brain disorders including epilepsy (32), ischemic stroke (33), traumatic brain injury (23), Alzheimer’s disease (34) and so on. Preventing seizure generation may have beneficial effects for curing these pathogenic conditions. It is well known that repeated seizures activate multiple cell death modes such as apoptosis (35). Targeting neuronal apoptosis was found to ameliorate the severity of seizures in experimental epilepsy models (36). Nevertheless, there is ample evidence supporting that treatment with the traditional anti-epileptic drugs via manipulating cell apoptosis induces many adverse reactions such as cognitive deficits and recurring seizures (37, 38). It is likely that seizure can initiate non-apoptotic forms of regulated cell death. Our recent work has delineated that ferroptosis, a novel cell death mode, is induced in FeCl_3_-, PTZ- and pilocarpine-induced seizure models (22, 23). Similarly, in our current study, we also provided direct evidence showing presence of ferroptosis in KA-induced epileptic seizure mice (small condensed mitochondria, elevations of 4-HNE and MDA and increased *Ptgs2* mRNA). And treatment with ferroptosis inhibitor Fer-1, Lipo or DFO dramatically attenuates epileptic seizures in this model, indicating a therapeutic potential of ferroptosis inhibition in treating seizure-associated disorders. Our data not only provides a previously unknown cell death mechanism suggesting that ferroptosis is activated in epileptic seizures, but also strongly implicates that reducing ferroptosis is a valid therapeutic target for seizure-related disorders such as epilepsy, ischemic stroke, traumatic brain injury and Alzheimer’s disease.

The molecular mechanism that underlies ferroptosis induction has also been investigated in our established KA-induced seizure mice model. We identify Lox as a novel ferroptotic regulator in neuronal toxicity induced by glutamate *in vitro* through gene expression profile analysis. Inhibition of Lox by genetic silencing or pharmacological treatment with BAPN suppresses ferroptotic cell death in hippocampal neurons *in vitro* and ameliorates KA-induced epileptic seizures *in vivo*.

The Lox family has five members including Lox and Loxl1-Loxl4. They are originally considered to be copper-dependent amino oxidases which are responsible for the cross-linking of collagen and elastin, thereby imparting the tensile strength and stability of tissues (24). Nowadays, there is substantial evidence supporting that Lox family possesses multiple biological functions, in addition to the intra- and inter-molecular cross-linking of collagen and elastin fibers. It has been shown that Lox promotes cancer migration, adhesion and metastasis in multiple types of cancers including colorectal cancer, breast cancer and lung cancer (26, 39, 40). Blockade of Lox by a pharmacological inhibitor BAPN or CCT365623 is able to delay tumor progression and cancer metastasis (40, 41). Moreover, the deleterious role of Lox in brain has been also observed in neuropathological states such as Alzheimer’s disease and spinal cord injury (42, 43). And pharmacological inhibition of Lox by BAPN has therapeutic effects in these disorders.

Likewise, our current data depicted elevation of Lox in both KA-induced epileptic seizures in mice and epileptic patients with hippocampal sclerosis (Figure 5A, Figure 5B and Figure 7A). And treatment with Lox inhibitor BAPN abrogated neuronal ferroptosis in vitro and in vivo. Other Lox family members including Loxl1, Loxl2, Loxl3 and Loxl4 were also detected in our present work and surprisingly these four isoforms were not altered in ferroptotic cell death model. These results make a strong emphasis on a key role of Lox in dictating hippocampal neuronal ferroptosis process following neurotoxicity. In fact, a previous investigation deciphered that Lox induced oxidative stress (an important factor for triggering ferroptosis) in vascular smooth muscle cells of spontaneously hypertensive rats, which supported, to some extent, an inducible role of Lox in ferroptosis process. Additionally, in this hypertensive model, Lox overexpression augmented oxidative stress that facilitated p38 MAPK activation. However, our present work revealed no significant effect of p38 after genetic silence or pharmacological inhibition of Lox following glutamate toxicity. One possible explanation for this discrepancy is attributable to different stressors. Interestingly, under glutamate oxidative toxicity conditions, ERK inactivation was found in our study after Lox inhibition. It was previously reported that ERK had dual roles, namely, protective or detrimental action, dependent on specific context. ERK may protect neurons against apoptosis but cause detrimental effects following necrotic death (30). In our glutamate-induced HT22 toxicity model, ferroptosis (a type of regulated necrosis) is a major sort of cell death mortality. We speculate that ERK activation is indispensible for induction of ferroptosis process. And our current work further proved that Lox activation triggered regulated necrosis, ferroptosis, in hippocampal neurons via ERK during glutamate stimuli. It was previously reported that ERK activation triggered Alox5 phosphorylation in leukocytes (44), which implicated that ERK-induced Alox5 activation served as a vital molecular mechanism in cellular biological process. Consistently, our present investigation also supported that elevation of phosphorylated Alox5 was found by ERK activation under glutamate- or erastin-induced ferroptosis in HT22 neurons. And inhibition of Alox5 activity by Zileuton evidently suppressed neuronal ferroptosis under glutamte or erastin exposure. Taken together, our findings delineate a Lox-ERK-Alox5 pathway as a novel signaling axis for inducing neuronal ferroptosis, which exacerbates seizure-induced hippocampal damage in epilepsy. We believe our data paves the way for future investigations on the therapeutic actions of Lox inhibitors in epilepsy or other seizure-associated disorders. Although Lox-based therapies may offer a new treatment strategy for refractory epilepsy, the potential side effects and toxicities of BAPN or new-generation Lox inhibitors must still be addressed. Future studies are essential to test the potential clinical implications of this therapeutic approach.

## Methods

### Establishment of KA-induced mouse seizure model and drug treatment

The KA-induced seizure model was established as previously described with minor modifications (22). Briefly, the mice were deeply anesthetized and mounted on a stereotaxic apparatus (RWD Life Science Co. Ltd., China). A 5 μl syringe (Hamilton, NV) was placed into the right dorsal hippocampus [anteroposterior (AP), −2 mm; mediolateral (ML), −1.8 mm; dorsoventral (DV), −2.3 mm] for KA injection (1.0 nmol of KA (Sigma-Aldrich, USA) in 1 μl of saline). After injection, the microsyringe was left in situ for additional 5 min to minimize the backflow along the injection trace. The control group was injected with equal volume of saline. Behavioral seizures were analyzed for the subsequent 6 h after KA injection according to a modified Racing scale (23, 45): stage 0, no response; stage 1, immobility; stage 2, rigidity; stage 3, head bobbing and circling; stage 4, intermittent rearing and falling; stage 5, continuous rearing and falling; and stage 6, tonic-clonic convulsions and rapid jumping. Animals that died were assigned stage 6 during the experiments. Prior to KA microinjection, BAPN, Fer-1 (Selleck, USA), Lipo-1 (Selleck, USA) and DFO (SantaCruz, USA) were administered intraperitoneally for 2 weeks, 2 weeks, 3 days and 1 week, respectively.

### Human brain tissue samples

Cortical tissues from drug-refractory TLE patients or brain trauma and hippocampal tissues from epileptic patients with hippocampal sclerosis were surgically obtained from XiangYa Hospital of Central South University.

### Ethical approval

All animal studies were handled according to the protocol approved by the Ethical Committee of the Animal Centre of Central South University. The study protocol related to human project were conducted in strict accordance with the Declaration of Helsinki and approved by the Ethics Committee of XiangYa Hospital of Central South University.

### Transmission electron microscopy

After reperfusion with 0.1 M phosphate buffer saline (PBS, pH=7.4), mice were killed and the brains from each group were carefully dissected. The brain samples were washed with precooled PBS. Targeted tissues were cut into 100 nm-thick sections. Selected brain areas were stained with uranyl acetate and lead citrate. Morphology of mitochondrial was viewed under a JEM2000EX transmission electron microscope (TEM; JEOL, Tokyo, Japan).

### Measurement of malonaldehyde (MDA) level

The relative concentration of MDA in cell or tissue lysates was measured using a commercial kit (S0131, Beyotime Technology Institute, China) according to manufacturer’s instructions.

### Real-time quantitative PCR

Total RNA was extracted using TRIzol reagent (15596018, Invitrogen, USA) following the manufacturer’s instructions. The procedure of real-time PCR was conducted as previously described (22). The detailed information of primer sequences used in our study was summarized in Table 1.

**Table 1.**
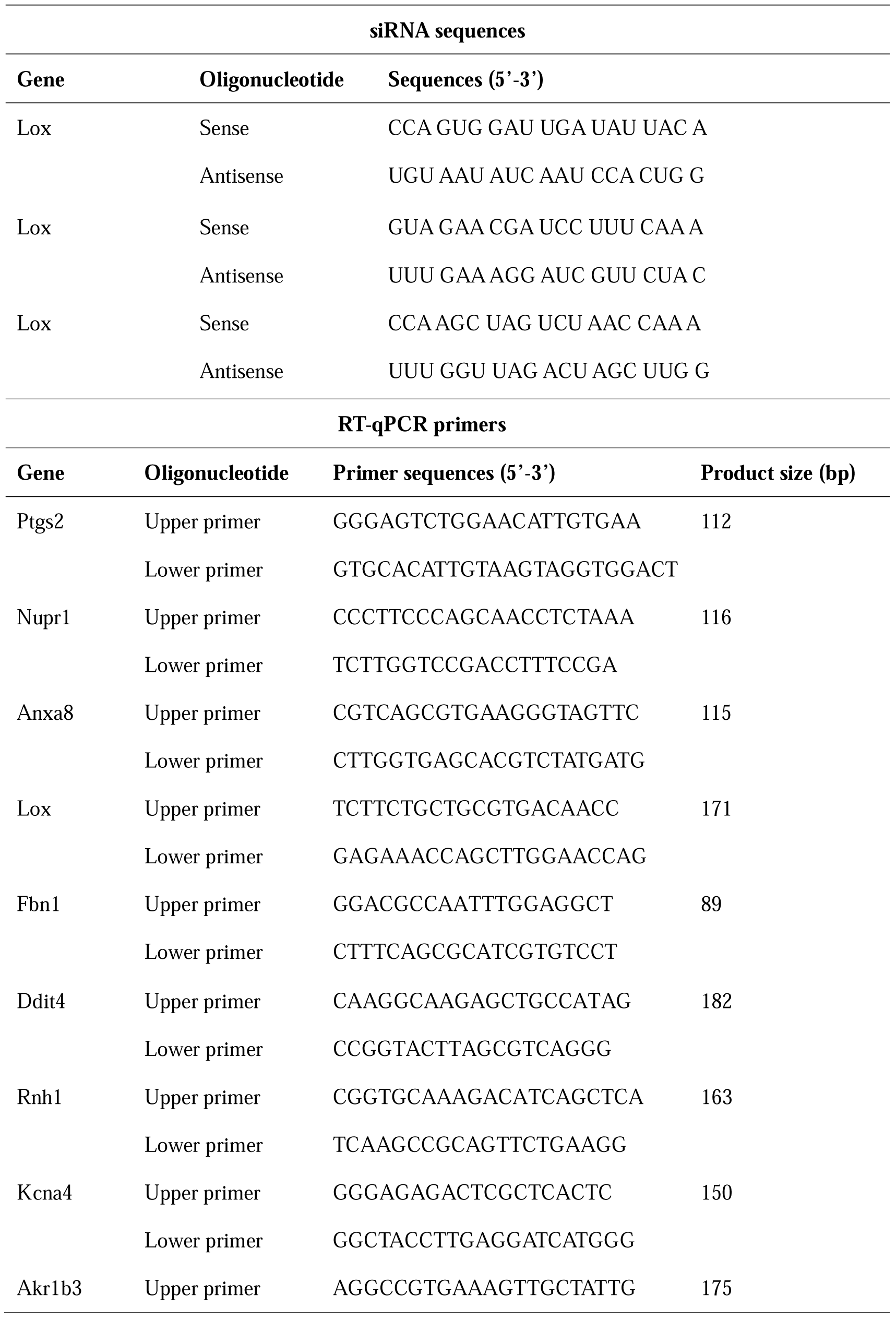
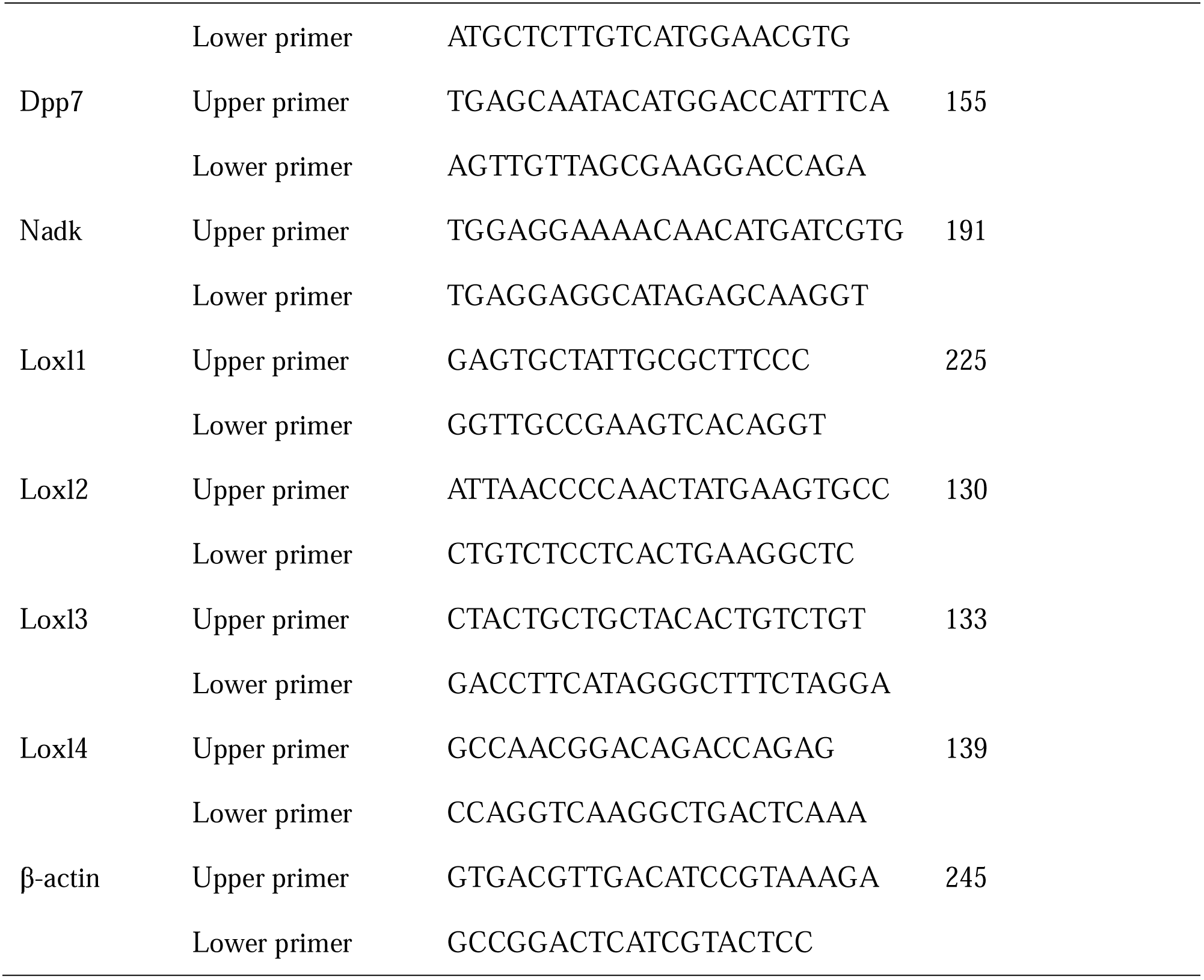
Oligonucleotide sequences used in the study.

### Western blotting assay

Immunoblot procedure was carried out according to our previous investigations (23). Briefly, *s*amples were lysed in protein lysis buffer (P0013, Beyotime Biotechnology Institue, China). Protein concentration was determined by a BCA protein assay kit (P0012, eyotime Technology Institute, China). Western blotting was carried out with primary antibodies including Lox, 4-HNE, ERK, p-ERK, JNK, p-JNK, p38, p-p38, Alox5, p-Alox5 Ser663, β-actin or α-tubulin. The detailed information of antibodies in our present work was listed in Table 2.

**Table 2.**
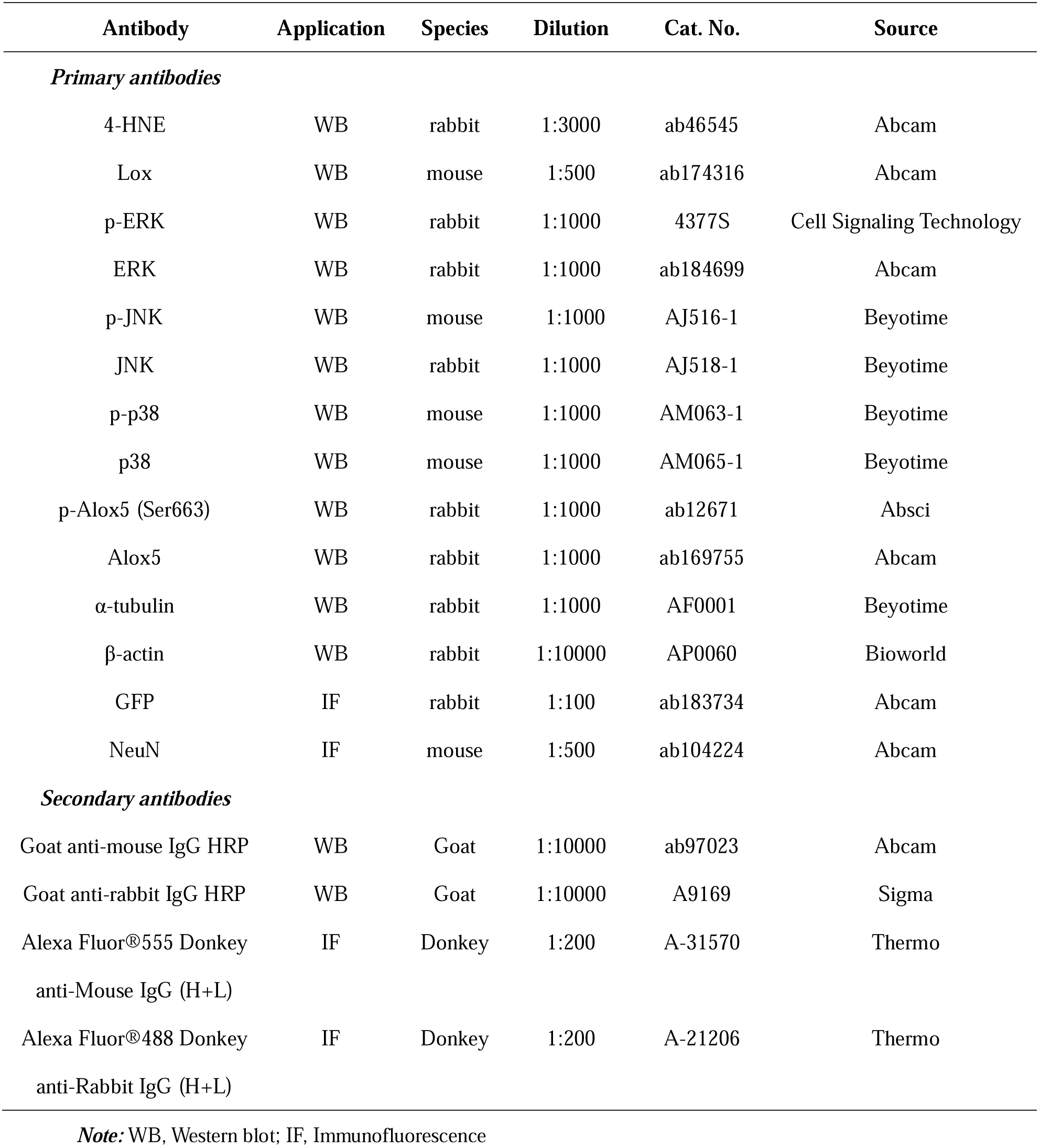
Antibodies used in the study.

### Cell culture

The immortalized hippocampal neuron HT22 was cultivated at 37 °C in a humidified 5% CO_2_ atmosphere and maintained in Dulbecco’s modified Eagle’s medium (DMEM, Gibco) with additions of 10% fetal bovine serum (FBS, Gibco), 100U/ml penicillin and 100 μg/ml streptomycin (Gibco).

### Gene expression profile

Total RNA from each group was extracted using TRIzol reagent (Invitrogen, USA). RNA quantification was performed by the NanoDrop ND-2000 (Thermo Scientific) and RNA integrity was evaluated using Agilent Bioanalyzer 2100 (Agilent Technologies). Gene expression profiling was conducted using Affymetrix MTA1.0 arrays (Affymetrix) following the manufacturer’s instructions. Differentially expressed genes were obtained with 2-fold change as well as a *p* value less than 0.05 between glutamate- and vehicle-treated groups.

### Cell death assessment by PI/Hoechst33342 staining

The HT22 cells were cultivated in 24-well plates at 20% density. After nearly 12 h, multiple drugs were added to the cell cultures for the indicated time, and then the neurons were stained with 5 μg/ml Propidium Iodide (PI) and 5 μg/ml Hoechst 33342 for 5 min at 37 °C. Cells from different groups were observed under a fluorescent microscope (Leica). Percentage of cell death was determined by PI/(PI +Hoechst 33342).

### CCK-8 assay

Cell viability was measured using a Cell Counting Kit-8 (CCK-8, Beyotime Biotechnology Institute, China). Overall, cells were incubated with 10 μl CCK-8 reagents in 100 μl growth medium for 2 h at 37°C. The absorbance for each sample was detected at 450 nm with a microplate reader. The results were repeated in triplicate.

### Measurement of Lox activity

Lox activity was determined using a commercial assay kit (ab112139, Abcam) according to manufacturer’s instructions.

### Analysis of lipid ROS by fluorescence-activated cell sorting (FACS)

Detection of cellular lipid ROS was performed as previously described with minor modifications (13). For short, HT22 cells were cultured in 6-well plates. After 12 h, cells were treated with test drugs for indicated times. Then HT22 neurons were harvested by trypsinizaiton, re-suspended in 500 μl Hanks Balanced Salt Solution (HBSS, Gibco, USA) containing 2 μM C11-BODIPY(581/591) (Thermo, USA) and placed at 37 °C in a cell culture incubator for 15 min. After one wash with HBSS, cells were re-suspended with fresh PBS and immediately analyzed by FACS using a flow cytometer (Beckman, USA) at an excitation wavelength of 488 nm. Data were obtained from FL1. A minimum of 20, 000 events were determined per replicate.

### RNA interference

All small interfering RNAs (siRNAs) were synthesized by RioBio (RioBio, China). The target siRNA sequences were summarized in Table 1. Before transfection, cells were seeded in 6-well or 96-well plates and grown to a confluence of about 40%. Lipofectamine™ RNAiMAX transfection reagent was employed following the manufacturer’s instructions. After 2 days of transfection, HT22 neurons were digested, passaged and cultivated in 24-well or 96-well plates, glutamate or erastin was then incubated for additional 12 h.

### Confocal microscopy

For preparation of brain tissue section, animals from each group were transcardially perfused with phosphate buffer saline (PBS) followed by 4% paraformaldehyde in 0.1 M phosphate buffer (PB). The whole brains were then carefully dissected and equilibrated in 30% sucrose in PB for over 3 days at 4 °C. Frozen tissue sections (8 µm) were collected using a cryostat (CM1900UV, Leica, Germany). For Confocal microscopic analysis, brain sections were washed in PBS for 10 min and then incubated with block solution (5% donkey serum in PBS and 0.2% Triton X-100) for 1 h at room temperature. Sections were then incubated overnight at 4°C with the primary antibody. After incubation, the slices were rinsed with PBS three times followed by incubation with the secondary antibody for 1 h. After washing, brain sections were counterstained with DAPI (1:500, Jackson Immunoresearch, USA) for 5 min at room temperature and mounted with anti-fading medium. Fluorescent images were obtained using a confocal laser scanning microscope (Nikon, Japan). To avoid the nonspecific staining, the incubation of primary antibody was replaced with normal serum, followed by all the subsequent experiments as described above. All the primary antibodies and secondary antibodies were summarized in Table 2.

### In vivo Lox overexpression by AAV vector

The AAV-mediated gene transfer method was used to cause Lox overexpression in the hippocampus under the control of SYN promoter. AAV-PHP.eB packing plasmid was provided by Obio Technology (Shanghai, China). Mice were anesthetized with 10% chloral hydrate before being fixed on a stereotaxic instrument (RWD, Shenzhen, China). For the AAV delivery, a microsyringe pump (RWD, Shenzhen, China) coupled with a 5 μl glass micropipette was used to stereotaxically inject 0.5 μl pAAV-SYN-EGFP-3FLAG (Virus titers: 0.75×1012 vg/ml) or 0.5 μl pAAV-SYN-Lox-P2A-EGFP-3FLAG (Virus titers: 1.04×1012 vg/ml) into the right hippocampus (anteroposterior [AP]: −2.00 mm; mediolateral [ML]: −1.8 mm; dorsoventral [DV]: −2.3 mm) at a rate of 4.6 nL/6 s. To determine the cell type-specific expression of AAV-PHP.eB-mediated gene transfer in the hippocampus, three animals per group were sacrificed for immunofluorescence staining at 4 wk after AAV-PHP.eB-GFP injection.

### Data analysis

All the results were displayed as mean ± SD. Comparison of 2 groups was performed using the dependent Student’s *t* test. For the comparison of more than 2 groups, we used ANOVA with the Bonferroni test. A *p* value of less than 0.05 was considered to be significant and was indicated \**p*<0.05, \*\**p*<0.01 or \*\*\**p*<0.001 in the graphs.

